# Realisation threshold revisited: does the minimum tooth size scale with body size?

**DOI:** 10.1101/2024.07.09.602691

**Authors:** Mona M. Christensen, Juha Laakkonen, Jukka Jernvall

## Abstract

Based on his analyses of lynx and brown bear teeth, Björn Kurtén coined the concept of ‘realisation threshold’, the smallest size at which a tooth can form and erupt properly. Kurtén found the smallest sizes of the studied lynx and bear teeth to be 2.5 mm and 3.5 mm, respectively, much larger than for example the smallest rodent teeth known to readily erupt. A recent study comparing developing teeth from shrews to elephants suggested a relatively unchanged theoretical minimum tooth size for mammalian teeth. Together, these studies have left open the question of whether realisation thresholds of teeth are larger in larger mammals than in small ones. In this study, we follow Kurtén’s line of thought and compare the sizes of teeth that are variably present in dentitions, and therefore likely cross the realisation threshold only occasionally. First, we show using published reports that variably present teeth are relatively small in large mammals, but larger than the previously suggested theoretical minimum tooth size. Next, we examine the canines of mares that are known to be variably present. We report one canine that, compared with information found in the literature, is by far the smallest compared to the body size. In conclusion, whereas the variably present teeth tend to be larger in large mammals, there may be overlooked potential for large mammals to develop very small teeth. This information can be helpful in extrapolating findings from common small model organisms, such as mice, to larger mammals, including humans.

## 1. Introduction

Loss of individual teeth and the emergence of supernumerary teeth are common in mammalian dentitions. In his doctoral thesis, Björn Kurtén (1953) discussed the relationship between tooth size and tooth emergence, analysing the mandibular third premolars (p3) of the brown bear (*Ursus arctos*), which in his dataset was present in 36% of the mandibles. Kurtén reported the greatest diameter of the smallest erupted p3 to be 3.5 mm, whereas it was 2.9 mm in two unerupted teeth. Kurtén suggested 3.5 mm to represent a “realisation threshold” for brown bear p3 emergence, a size under which this tooth would not correctly form or erupt.

In addition to bears, Kurtén (1963) described a case of evolutionary tooth re-emergence in the lynx (*Lynx lynx*). The mandibular second molar (m2) was lost in known Felidae in the Miocene but was present in approximately 10% of lynx dentitions studied. Kurtén found the smallest lynx m2 to be 2.5 mm in width and length, and proposed that the occasional re-appearance may be related to m2 crossing its realisation threshold size. Together, Kurtén’s studies on bears and lynxes suggest that tooth size may influence tooth emergence. The realisation threshold is also an example of Kurtén’s work with both evolutionary and developmental implications that has attracted continuing research interest (e.g. Lynch 2022, Christensen *et al*. 2023).

In addition to animals, tooth size has been linked to variation in tooth number in humans. Brook (1984) proposed a model where the probability for tooth loss increases with small tooth size, and probability for supernumerary tooth emergence increases with large tooth size (Brook 1984). Clinical data support this model by showing that individuals with fewer teeth than usual tend to have smaller average tooth size compared to healthy, control group individuals (Baum & Cohen 1971, Schalk-Van der Weide *et al*. 1994, and Brook *et al*. 2009). In contrast, individuals with supernumerary teeth tend to have a large average tooth size (Khalaf *et al*. 2005 and Brook *et al*. 2009). Additionally, the teeth of family members of hypodontia patients (defined as lacking more than six teeth congenitally) tend to be smaller than those of control group individuals (Schalk-Van der Weide & Bosman 1996, McKeown 2002). Brook’s model can be considered analogous to Kurtén’s theory of realisation threshold in that it connects tooth size with tooth loss and emergence.

Kurtén’s realisation threshold appears to have been partially inspired by Grüneberg’s work on mouse teeth. Based on measurements of mouse teeth, Grüneberg (1951) suggested that during development, the probability for a functional tooth to form decreases if the tooth germ size falls below a certain threshold. During the past decades this threshold has garnered cumulative support from works on the developmental genetics of mouse molars. These studies have revealed several keystone genes required for the progression of tooth development beyond the cap stage (Hallikas *et al*. 2021). At this developmental stage the epithelial cervical loops grow laterally and start to encompass the underlying mesenchyme to form the crown. This process is regulated by an epithelial signalling centre, the primary enamel knot, which expresses many of the keystone genes of crown formation (Hallikas *et al*. 2021).

Mouse teeth are miniscule in size compared that of the bear or lynx. This raises the question whether the realisation threshold scales with the tooth size. Patterning of tooth cusps happens in larger size in larger teeth (Jernvall 1995, Christensen *et al*. 2023), which has led to a proposal that realisation threshold may scale with the final tooth size (Jernvall 1995). This scaling of patterning has been recently shown to result from the integration of growth and patterning by insulin-like growth factor (IGF) signalling (Christensen *et al*. 2023). In contrast to the scaling of the patterning, however, Christensen *et al*. (2023) also showed that cap stage tooth germs are largely similar in width across mammals. Thus, assuming that the realisation threshold is linked to the emergence of the cap stage during tooth development, we might expect even large species to be able to initiate tooth development in a small size.

Here we address the scaling of realisation thresholds by combining evidence from the literature with empirical data from a large mammal, the horse (*Equus caballus*). Although horses have generally large teeth, the canines of mares (female horses) are small, and often missing (Anthony *et al*. 2010). Stallions and geldings (male horses), in contrast, have typically relatively large canines. We follow Kurtén’s line of thought and focus on teeth that are variably present in a species. We define variably present teeth as those that are not expected based on the normal dental pattern of the taxa (supernumerary teeth) and those that are included in the dental pattern but are present at low frequencies (such as the mare canines). The rationale behind this approach is that variably present teeth only occasionally cross the realisation threshold, and therefore can be expected to be near the lower limit in size for each species.

## 2. Material and methods

### 2.1. Analyses of published cases

A literature search was conducted to find size data on variably present teeth. For each species with reported variably present teeth, we tabulated head and body length, variably present tooth size and shape and first mandibular molar size (Table 1). All body and head length data were taken from Nowak (1999), where minimum and maximum lengths are reported, and the average of these was calculated. Bucco-lingual dimensions of the variably present teeth were tabulated, except in the case of *U. arctos* and elk *(Alces alces*) where the measurement was reported to be the greatest crown diameter (Kurtén 1953, Steele & Parama 1979) and for roe deer (*Capreolus capreolus*) where the tooth orientation was not certain (Chaplin & Atkinson 1968). In these cases, the tooth shape was conical, so the measurements were accepted as proxies for the bucco-lingual width. When several specimens of the same tooth were available, the smallest one of these was used for the analysis, as this was reasoned to be closest to the realisation threshold size for the particular tooth. Tooth shapes were based on the author’s description or estimated visually when possible.

**Table 1.**
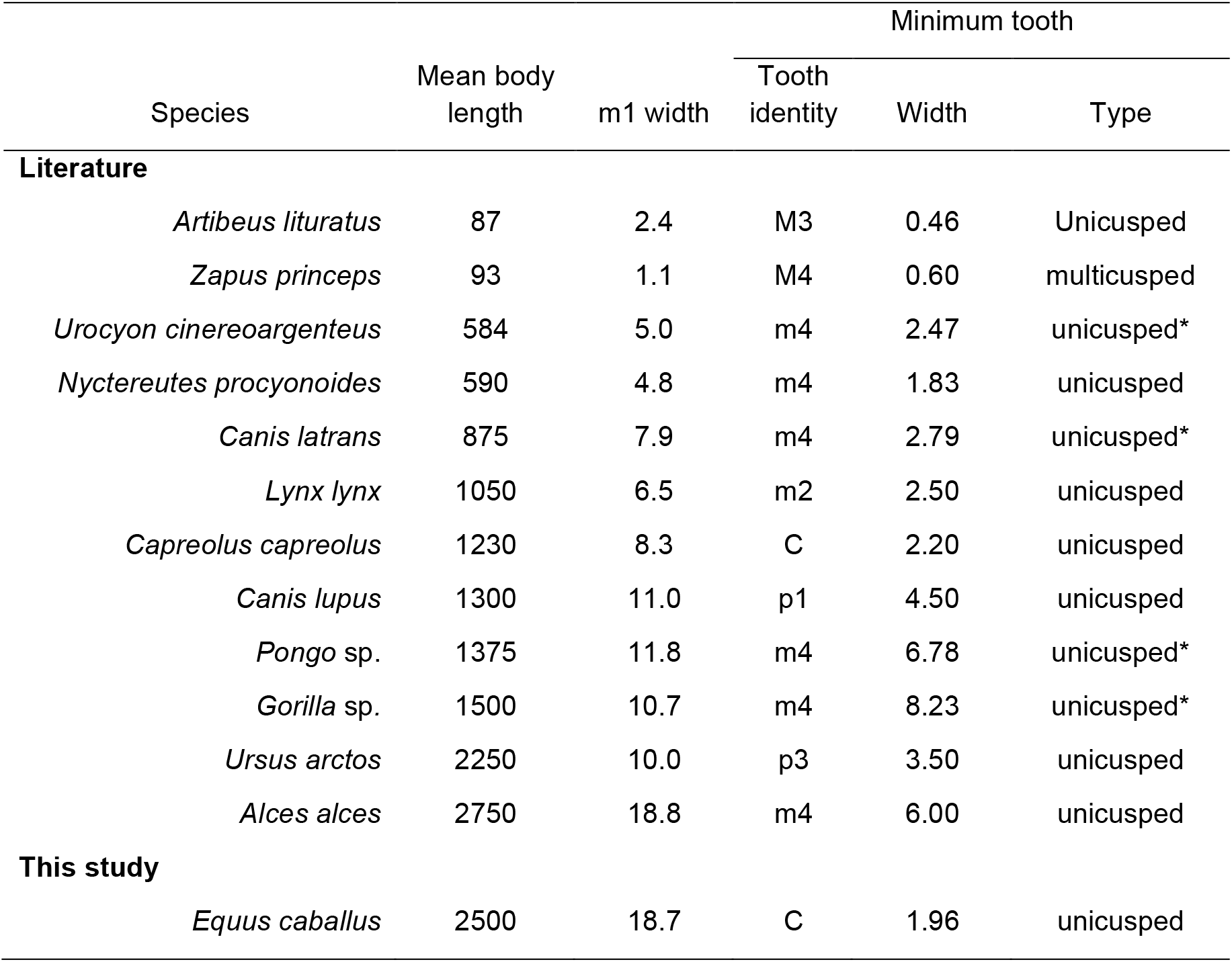
Tooth sizes for supernumerous or variably present teeth in mammals of different body size. Upper and lower case in in tooth identity denotes to maxillary and mandibular teeth, respectively. Units are in millimetres. Asterisk in unicusped tooth type indicates a tooth with a cingulid.

| Species | Mean body length | m1 width | Minimum tooth |  |  |
| --- | --- | --- | --- | --- | --- |
|  |  |  | Tooth identity | Width | Type |
| Literature |  |  |  |  |  |
| *Artibeus lituratus* | 87 | 2.4 | M3 | 0.46 | Unicusped |
| *Zapus princeps* | 93 | 1.1 | M4 | 0.60 | multicusped |
| *Urocyon cinereoargenteus* | 584 | 5.0 | m4 | 2.47 | unicusped* |
| *Nyctereutes procyonoides* | 590 | 4.8 | m4 | 1.83 | unicusped |
| *Canis latrans* | 875 | 7.9 | m4 | 2.79 | unicusped* |
| *Lynx lynx* | 1050 | 6.5 | m2 | 2.50 | unicusped |
| *Capreolus capreolus* | 1230 | 8.3 | C | 2.20 | unicusped |
| *Canis lupus* | 1300 | 11.0 | p1 | 4.50 | unicusped |
| *Pongo* sp. | 1375 | 11.8 | m4 | 6.78 | unicusped* |
| *Gorilla* sp. | 1500 | 10.7 | m4 | 8.23 | unicusped* |
| *Ursus arctos* | 2250 | 10.0 | p3 | 3.50 | unicusped |
| *Alces alces* | 2750 | 18.8 | m4 | 6.00 | unicusped |
| This study |  |  |  |  |  |
| *Equus caballus* | 2500 | 18.7 | C | 1.96 | unicusped |

Because postcanine tooth size generally scales with body size (Fortelius 1985, Copes & Schwartz 2010), and the first molar is the least variable of molars in size (Gingerich 1974), we used the first mandibular molar as a control to which we compared the scaling of the variably present teeth.

The tooth data were obtained for the following species, either from text or measured from the figures, coyote (*Canis latrans*), gray fox (*Urocyuon cinereoargenteus*), and raccoon dog (*Nyctereutes procyonoides*) (Asahara 2016), gorilla (*Gorilla* sp.) and orangutan (*Pongo* sp.) (Schwartz 1984), jumping mouse (*Zapus princeps*) (Krutzsch 1953, Klineger 1963), brown bear (*Ursus arctos*) (Kurtén 1953, Stenberg *et al*. 2024), lynx (Kurtén 1963, Gomerčić *et al*. 2010), roe deer (*Capreolus capreolus*) (Chaplin & Atkinson 1968, Chirichella *et al*. 2021), great fruit-eating bat (*Artibeus lituratus*) (Rui and Drehmer 2005), gray wolf (*Canis lupus*) (Buchalczyk *et al*. 1980, Baryshnikov *et al*. 2009), and elk (*Alces alces*) (Steele & Parama 1979, Pasda *et al*. 2020).

### 2.2. Horse canines

Horse material was donated by the owners for teaching and research at the Veterinary Faculty of the University of Helsinki and consisted of various breeds but only adult individuals. The heads of all specimens were examined for supernumerary teeth and other dental anomalies. Of the nineteen mares none showed erupted canines, five had prominence of the gum in the mandible, of which three covered partially developed canines (the canines were extracted with a bone-saw). Maxillary diastema lacked visible development of teeth, but closer examination of one skull revealed a rudimentary canine that was by far the smallest of the recovered teeth. This skull was cleaned and the tooth was μCT scanned using Bruker 1272 μCT scanner following (Christensen *et al*. 2023). Image preparation were done as in (Christensen *et al*. 2023) and enamel thickness measurements using Hausdorff distance in Meshlab (version 2022.5). The voxel size was 13 μm.

## 3. Results

### 3.1. Minimum tooth size shows a slight trend to increase with body size

A plot using the mandibular first molar (m1) width shows how large species have larger molars than small species (Fig. 1A, using the units in the figure, the linear regression slope is 5.99 and the intercept is 1.524, *r*^2^ = 0.88). In contrast, the variably present teeth are smaller but there is a great deal of variation in the published cases. Teeth with more than one cusp, or a cingulid surrounding a single cusp, appear to be larger in larger species. Especially the extra molars of the gorilla and orangutan are relatively large, well over half the width of the anterior molars [Fig. 1B, Table 1, Schwartz (1984)]. The linear regression slope is 5.33 and the intercept is -0.546 (*r*^2^ = 0.93). To the extent that cingulids can be considered to reflect development of additional cusps, the increase agrees with multiple cusps requiring larger growth domain to appear during development (Harjunmaa *et al*. 2014, Christensen *et al*. 2023). Examining single cusped teeth only, they show comparatively little increase in larger species, but this is affected by the inclusion of the mare canine in data (Fig. 1B). Excluding the mare, the linear regression slope is 1.77 and the intercept is 0.661 (*r*^2^ = 0.78). Including the mare, the linear regression slope is 1.25 and the intercept is 1.026 (*r*^2^ = 0.47). The width of the mare canine is close to the extra molar of the raccoon dog (*Nyctereutes*, Table 1), a much smaller species.

**Fig. 1.**
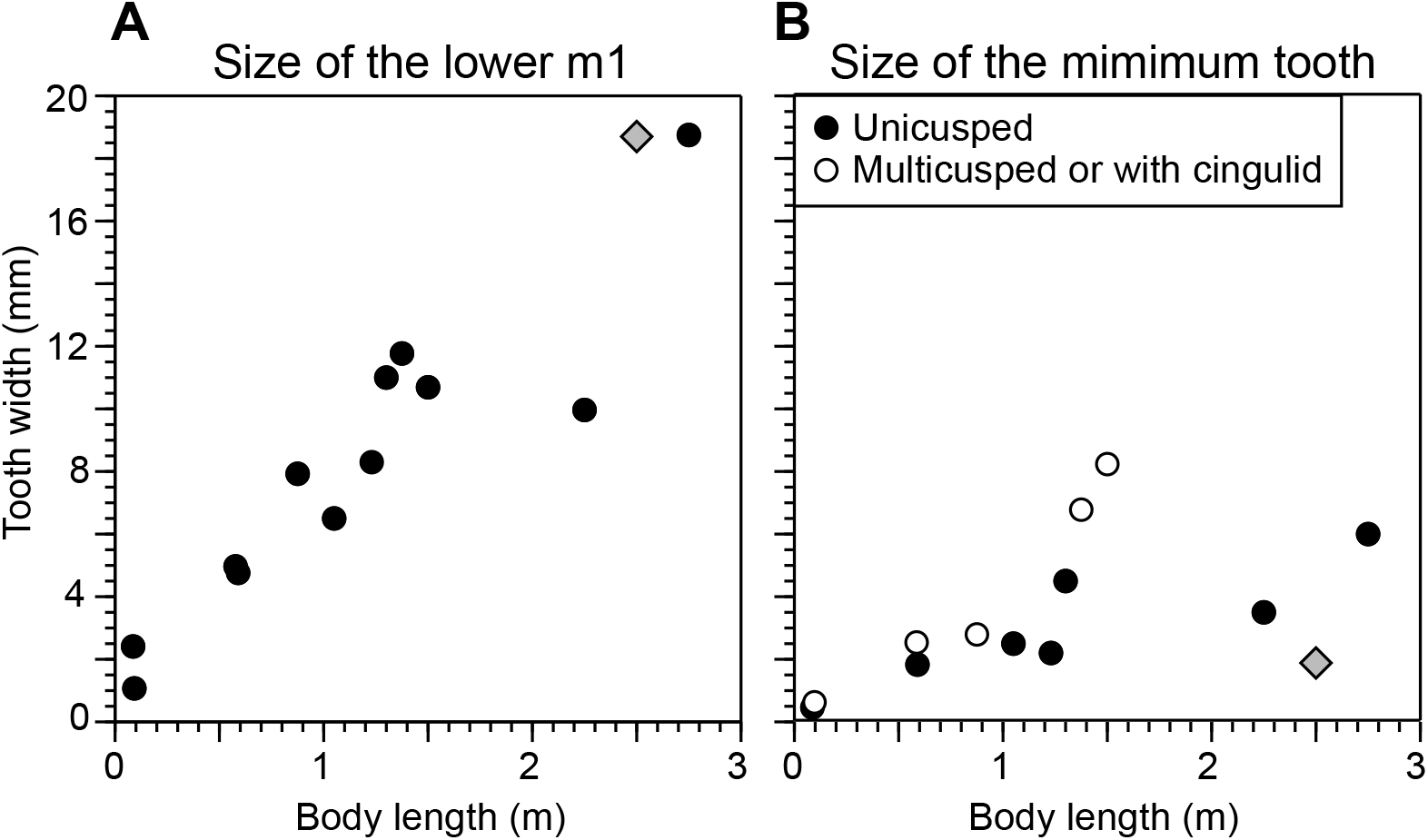
Body length compared with widths of the mandibular first molar (m1) and variably present teeth. (**A**) Whereas the m1 size increases with the body size, even when comparing diverse taxonomic groups (Table 1), (**B**) the variably present teeth are much smaller and show a weaker association with the body size. The diamond marks the horse measurements, and in (**B**) the mare canine studied in this paper.

### 3.2. The canine of the mare illustrates how a smallest possible tooth is formed in a large mammal

The mare with the left maxillary canine had both mandibular canines present but unerupted. The socket for right maxillary canine was present, but the tooth had been lost either in life or during preparation. The left maxillary canine was in the socket, but was only detectable after cleaning of the skull (Fig. 2). Compared to the left maxillary socket and canine, the socket for the right canine was larger, and the unerupted mandibular canines were visibly larger. From this we infer that the left maxillary canine is the smallest tooth present in this skull (Fig. 2). The canine has a long root and round peg-shaped crown. The overall length from the tip to the end of the root is 16.2 mm (Fig. 3A). The root is slightly curved, having a round cross section at the neck (Fig. 3B), and gradually flattening towards the end of the root. The pulpal cavity is almost completely ossified, becoming visible 1.3 mm below the crown, and roughly 60 μm wide (Fig. 3B). The root is covered with a thick layer of cementum ranging from 0.3 to 0.5 mm in thickness for most parts of the root (Fig. 3A, B). Closer to the crown, there are large areas where the cementum has been lost, possibly due to resorption in life. Portions of the enamel close to the tooth neck are covered by a thin layer of cementum. The neck, excluding the cementum layer, is 1.4 to 1.5 mm wide.

**Fig 2.**
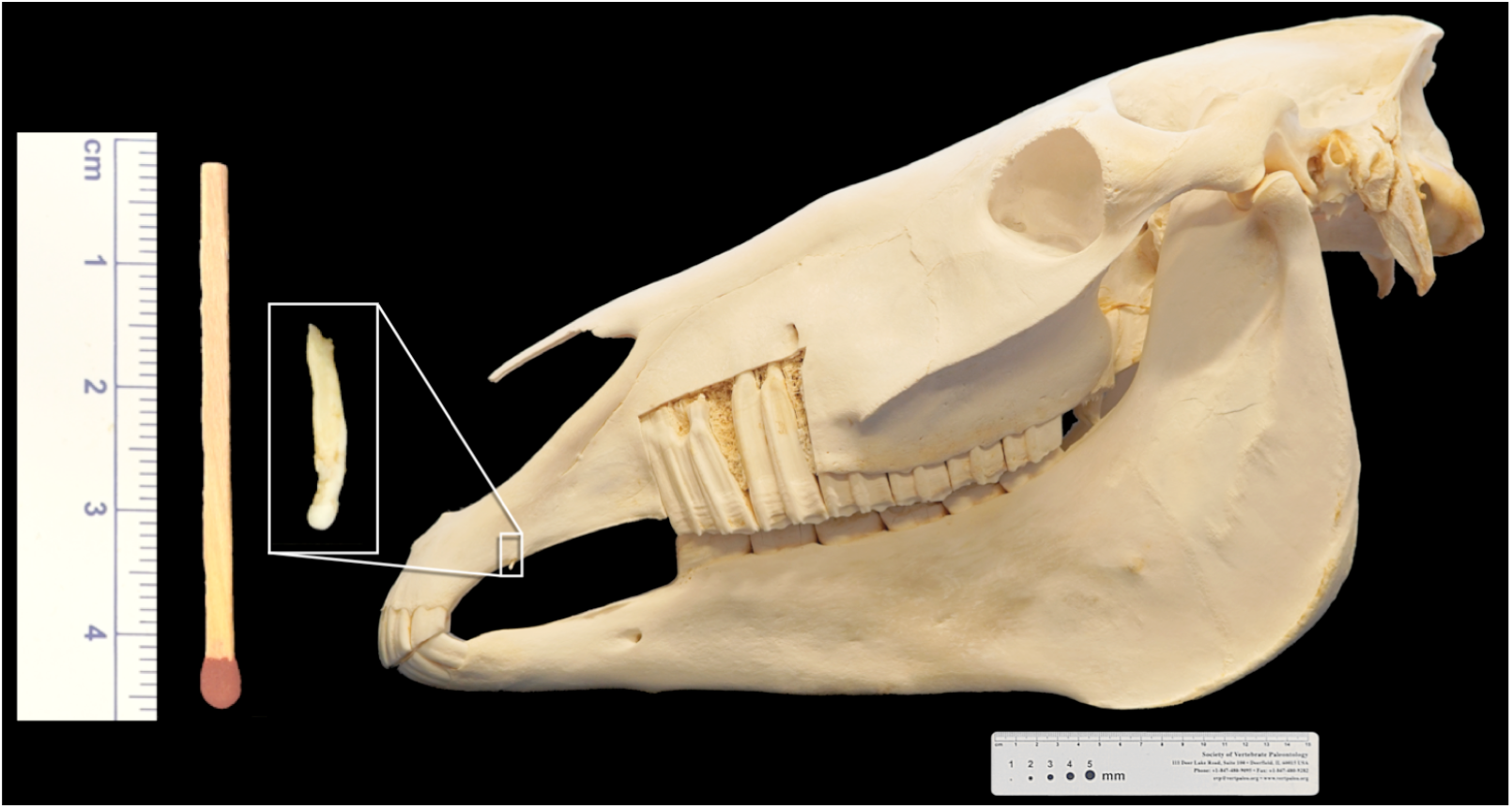
A mare skull with extracted left maxillary canine shown on the left, with a matchstick for a scale. Two maxillary premolars are exposed to illustrate the size of the postcanine teeth.

**Fig 3.**
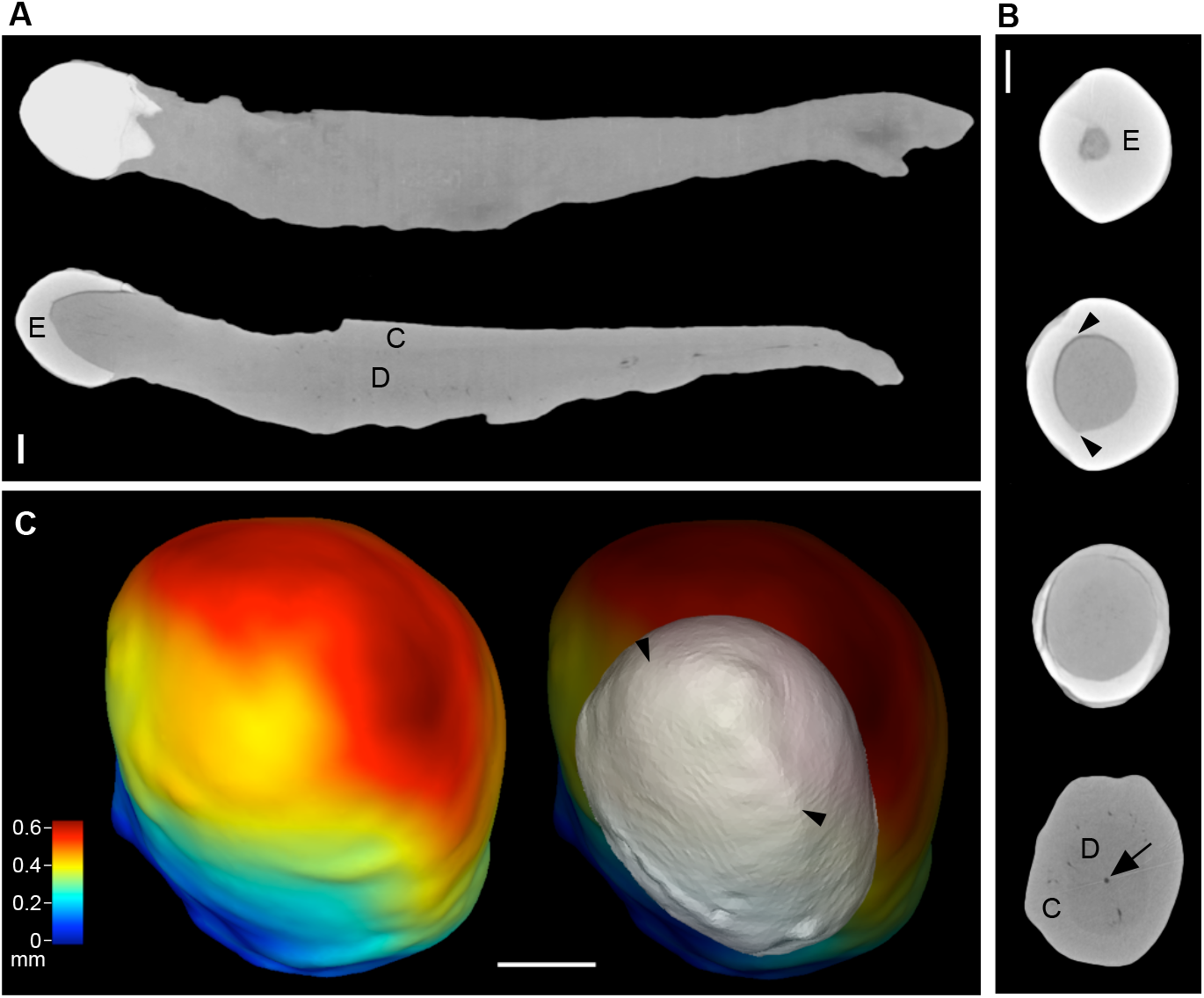
Morphological analyses of the mare canine (**A)** A μCT reconstruction of the canine showing a surface image and mid-section. (**B**) Cross sections near the tip showing the dentine tip (top), mid crown section showing dentine ridges (arrowheads), a section from the neck, and from the middle of the root where the narrow pulp cavity is still visible (arrow). (**C**) Three-dimensional reconstruction showing the enamel thickness as a heatmap, and the underlying EDJ in grey with the lateral ridges (arrowheads). Enamel (E), dentine (D), cementum (C), Scale bars, 0.5 mm.

The crown has the overall appearance of a round peg being 2.1 mm high (Fig. 3). However, a closer examinations shows that the cross section is slightly flattened (Fig. 3B, C). There is a rounder side with thicker enamel, and more flat side with thinner enamel. These sides are divided by lateral ridges visible at the enamel-dentine junction (EDJ) and by thicker enamel above the ridges (Fig. 3B, C). Together these morphological features suggest an incipient canine morphology. At the tip of the EDJ (Fig. 3C, top), the ridges are no longer visible and the width of the dentine tip is around 0.3 to 0.35 mm, before ending altogether about 0.05 mm further up. Dentinal tubules can be detected almost to the tip of the EDJ, suggesting this to be the origination point of the crown development.

## 4. Discussion

Overall, when examining several species, teeth that are variably present are, when present, relatively small. The ‘relatively small’ is the topic that is brought now into focus be the study of Christensen *et al*. (2023). Collecting histological data from 13 species, ranging in size from the shrew (*Sorex araneus*) to elephant (*Loxodonta africana*), the early cap stage tooth width was found to range from 0.15 to 0.21 mm. There was very little increase in size of the cap stage in large teeth across this body size range. From a regression slope fitted to the data, one and ten millimetre wide teeth would have cap stages that are 0.17 and 0.18 mm wide, respectively. Basically this is the difference between mouse and human molars. Even broader teeth such rhinoceros molars (Fortelius 1985) reaching 50 mm in width are predicted to have cap stages that are only 0.20 mm wide. These cap stage sizes were proposed by Christensen *et al*. (2023) to be close to the lower theoretical size limit for mammalian teeth. This prediction seems to be supported by teeth of genetically modified mice and small bodied fossil taxa (Luo *et al*. 2022, Christensen *et al*. 2023). Because the observed minimum tooth sizes approaching the theoretical limit were all from small mammals, the question of the minimum tooth size in larger mammals remains. Hence, does Kurtén’s realisation threshold scale with the body size?

The examples obtained from the literature show that variably present teeth are small compared to their ‘normal’ counterparts (Fig. 1). Yet, these teeth are larger than would be expected from the cap stage expectation (Christensen *et al*. 2023). There is also a tendency for larger mammals to have larger variably present teeth, even when they have only one cusp (Fig. 1B). One reason for the increase of the minimum tooth size in larger species could be smaller teeth becoming more easily reabsorbed, perhaps because they simply fail to erupt through the thicker oral epithelium. Kurtén (1953) already considered failure to erupt as a possible reason for the drop in the frequency of the smallest premolars in bears.

Another reason for the lack of tiny teeth in large mammals could be sampling. Very small teeth are easily overlooked, and lost during preparation of specimens. This is the reason why we examined the canines of the mares. For a sample of 297 mares, Anthony *et al*. (2010) reported that 24.6% had canines and only 8.1% had all four of them. In our examinations, fully formed canines were found in 21% of the mares (4/19). One of these was likely to have had all four canines, including the small maxillary canine studied here (Fig. 2). This tooth is exceptionally small for the body size and rest of the teeth (Figs 1B, 2). Because our comparative measurements included enamel (Fig. 1), they overestimate the underlying dentine. Thus, EDJ size corresponds to the developmental size more accurately. Examining the mare canine EDJ size, the cusp tip seems to be approaching the theoretical lower limit for tooth size by having width of only 0.3 mm (Fig. 3B, C). Although such a specimen can be considered a lucky discovery, its mere existence implies for the potential for the realisation threshold to be relatively body size invariant. In line with this are the reportedly smallest incisors and premolars found in humans affected by by microcephalic osteodysplastic primordial dwarfism type II (MOPD II). These teeth measured 2 to 2.5 mm in width (Kantaputra *et al*. 2011).

In conclusion, whereas the variably present teeth tend to be larger in large mammals than in small ones, the developmental potential for large mammals to produce very small teeth remains. To decipher the actual limits for minimum tooth size requires thorough sampling, and careful preparation of specimens. With modern imaging techniques, it should be possible to obtain better appraisals of the smallest teeth in the largest mammals. Although search for the smallest teeth might seem like an academic exercise, it does test our theories about development. Currently significant effort is invested in regenerative medicine. Tissue and organ engineering combined with stem cell biology aims to provide tools to regenerate organs such as teeth. Much of this knowledge is based on work carried out using mice. Part of extrapolating the knowledge from mice to large mammals is examining whether and to what extent large mammals can scale downwards.

## Acknowledgments

We thank Aida Kaffash Hoshiar for μCT scanning and the members of Jernvall lab for comments and discussions on this work. This study was supported by the Sigrid Jusélius Foundation (J.J.) and Doctoral Programme in Biomedicine (M.M.C.).

## Notes

### Competing Interest Statement

The authors have declared no competing interest.

## References

Anthony, J., Waldner, C., Grier, C., & Laycock, A. R. 2010: A Survey of Equine Oral Pathology. ---Journal of Veterinary Dentistry 27: 12–15.

Asahara, M. 2016: The origin of the lower fourth molar in canids, inferred by individual variation. ---PeerJ 4:e2689, doi: 10.7717/peerj.2689.

Baryshnikov, G. F., Mol, D., & Tkhomov, A. N. 2009: Finding of the Late Pleistocene carnivores in Taimyr Peninsula (Russia, Siberia) with paleoecological context. ---Russian Journal of Theriology 8: 107–113.

Baum, B. J., & Cohen, M. M. 1971: Agenesis and tooth size in the permanent dentition. ---Angle Orthodontist 41: 100–102.

Brook, A. H. 1984: A unifying aetiological explanation for anomalies of human tooth number and size. ---Archives of Oral Biology 29: 373–378.

Brook, A. H., Griffin, R. C., Smith, R. N., Townsend, G. C., Kaur, G., Davis, G. R. & Fearne, J. 2009: Tooth size patterns in patients with hypodontia and supernumerary teeth. ---Archives of Oral Biology 54 Supplement 1: S63–S70.

Buchalczyk, T., Dynowski, J. & Szteyn, S. 1981: Variations in number of teeth and asymmetry of the skull in the wolf. ---Acta Theriologica 26: 23–30.

Chaplin, R. E., & Atkinson, J. 1968: The occurrence of upper canine teeth in Roe deer (Capreolus capreolus) from England and Scotland. ---Journal of Zoology 155: 141–144.

Chirichella, R., Marinis, A. M. D., Pokorny, B. & Apollonio, M. 2021: Dentition and body condition: tooth wear as a correlate of weight loss in roe deer. ---Frontiers in Zoology 18: 47, doi: 10.1186/s12983-021-00433-w.

Christensen, M. M., Hallikas, O., Das Roy, R., Väänänen, V., Stenberg, O. E., Häkkinen, T. J., François, J-C., Asher, R. J., Klein, O. D., Holzenberger, M. & Jernvall, J. 2023: The developmental basis for scaling of mammalian tooth size. ---Proceedings of the National Academy of Sciences U.S.A. 120:e2300374120, doi: 10.1073/pnas.2300374120.

Copes, L. E. & Schwartz, G. T. 2010: The scale of it all: postcanine tooth size, the taxon-level effect, and the universality of Gould’s scaling law. ---Paleobiology 36: 188–203.

Fortelius, M. 1985: Ungulate cheek teeth: developmental, functional, and evolutionary interrelations. ---Acta Zoologica Fennica 180: 1–76.

Gingerich, P. D. 1974: Size Variability of the Teeth in Living Mammals and the Diagnosis of Closely Related Sympatric Fossil Species. ---Journal of Paleontology 48: 895–903.

Gomerčić, T., Sindičić, M., Gomerčić, M.Đ., Gužvica, G., Frković A., Pavlović, D., Kusak, J., Galov, A. & Huber, Đ. 2010: Cranial morphometry of the Eurasian lynx (Lynx lynx L.) from Croatia. ---Veterinarski Arhiv, 80: 393–410.

Grüneberg, H. 1951: The genetics of a tooth defect in the mouse. ---Proceedings of the Royal Society of London. Series B - Biological Sciences 138: 437–451.

Hallikas, O., Das Roy, R., Christensen, M. M., Renvoisé, E., Sulic, A. & Jernvall, J. 2021: System-level analyses of keystone genes required for mammalian tooth development. ---Journal of Experimental Zoology Part B: Molecular and Developmental Evolution 336: 7–17.

Harjunmaa, E., Seidel, K., Häkkinen, T., Renvoisé, E., Corfe, I. J., Kallonen, A., Zhang, Z-Q., Evans, A. R., Salazar-Ciudad, I., Klein, O. & Jernvall, J. 2014: Replaying evolutionary transitions from the dental fossil record. ---Nature 512: 44–48.

Jernvall, J. 1995: Mammalian molar cusp patterns: Developmental mechanisms of diversity. ---Acta Zoologica Fennica 198: 1–61.

Kantaputra, P., Tanpaiboon, P., Porntaveetus, T., Ohazama, A., Sharpe, P., Rauch, A., Hussadaloy, A. & Thiel, C. T. 2011: The smallest teeth in the world are caused by mutations in the PCNT gene. ---American Journal of Medical Genetics Part A 155: 1398–1403.

Khalaf, K., Robinson, D. L., Elcock, C., Smith, R. N. & Brook, A. H. 2005: Tooth size in patients with supernumerary teeth and a control group measured by image analysis system. ---Archives of Oral Biology 50: 243–248.

Klingener, D. 1963: Dental Evolution of Zapus. ---Journal of Mammalogy 44: 248–260.

Krutzsch, P. 1953: Supernumerary molars in the jumping mouse (Zapus princeps). ---Journal of Mammalogy 34: 265–266.

Kurtén, B. 1953: On the variation and population dynamics of fossil and recent mammal populations. ---Acta Zoologica Fennica 76:1–122.

Kurtén, B. 1963: Return of a lost structure in the evolution of the felid dentition. ---Societas Scientarium Fennica Commentationes Biologicae 26:1–12.

Luo, Z., Bhullar, B.-A., Crompton, A., Neander, A. & Rowe, T. 2022: Reexamination of the Mandibular and Dental Morphology of the Early Jurassic Mammaliaform Hadrocodium wui. ---Acta Palaeontologica Polonica 67: 95–113.

Lynch, V. 2022: Is there a loophole in Dollo’s law? A DevoEvo perspective on irreversibility (of felid dentition). ---Journal of Experimental Zoology Part B: Molecular and Developmental Evolution 340: 509–517.

McKeown, H. F., Robinson, D. L., Elcock, C., Al-Sharood, M. & Brook, A. H. 2002: Tooth dimensions in hypodontia patients, their unaffected relatives and a control group measured by a new image analysis system. ---The European Journal of Orthodontics 24: 131—141.

Nowak, R. M. 1999: Walker’s Mammals of the World (6th ed.), vol. 1. ---The Johns Hopkins University Press, Baltimore.

Pasda, K., Correa, M. L., Stojakowits, P., Häck, B., Prieto, J., al-Fudhaili, N. & Mayr C. 2020: Cave finds indicate elk (Alces alces) hunting during the Late Iron Age in the Bavarian Alps.---E&G Quaternary Science Journal 69: 187–200.

Rui, A. M. & Drehmer, C. J. 2004: Anomalies and variation in the dental formula of bats of the genus Artibeus Leach (Chiroptera, Phyllostomidae). ---Revista Brasileira de Zoologia 21: 639—648. [in Portugese with English abstract]

Schalk-Van der Weide, Y., Steen, W. H. A., Beemer, F. A. & Bosman, F. 1994: Reductions in size and left-right asymmetry of teeth in human oligodontia. ---Archives of Oral Biology 39: 935–939.

Schalk-Van der Weide, Y. & Bosman, F. 1996: Tooth size in relatives of individuals with oligodontia. ---Archives of Oral Biology 41: 469–472.

Schwartz, J. H. 1984: Supernumerary teeth in anthropoid primates and models of tooth development. ---Archives of Oral Biology 29: 833–842.

Steele, D. G. & Parama, W. D. 1979: Supernumerary Teeth in Moose and Variations in Tooth Number in North American Cervidae. ---Journal of Mammalogy 60: 852–854.

Stenberg, O. E., Moustakas-Verho, J. E. & Jernvall, J. 2024: Rewinding developmental trajectories shows how bears break a developmental rule. ---Annales Zoologici Fennici 61: in press.

